# APOE regulates the transport of GM1

**DOI:** 10.1101/2024.04.02.587789

**Authors:** Dong Yan Zhang, Jian Wang, Gangtong Huang, Sara Langberg, Feng Ding, Nikolay V. Dokholyan

**Author notes:** Corresponding author: Nikolay V. Dokholyan,.

## Abstract

Apolipoprotein E (APOE) is responsible for lipid transport, including cholesterol transport and clearance. While the ε4 allele of APOE (APOE4) is associated with a significant genetic risk factor for late-onset Alzheimer’s disease (AD), no mechanistic understanding of its contribution to AD etiology has been established yet. In addition to cholesterol, monosialotetrahexosylganglioside (GM1) is a crucial lipid component in cell membranes and has been implicated in promoting the aggregation of amyloid beta protein (Aβ), a key protein associated with AD. Here, we ask whether there are direct interactions between APOE and GM1 that further impact AD pathology. We find that both APOE3 and APOE4 exhibit superior binding affinity to GM1 compared to cholesterol and have an enhanced cellular uptake to GM1 lipid structures than cholesterol lipid structures. APOE regulates the transport process of GM1 depending on the cell type, which is influenced by the expression of APOE receptors in different cell lines and alters GM1 contents in cell membranes. We also find that the presence of GM1 alters the secondary structure of APOE3 and APOE4 and enhances the binding affinity between APOE and its receptor low-density lipoprotein receptor (LDLR), consequently promoting the cellular uptake of lipid structures in the presence of APOE. To understand the enhanced cellular uptake observed in lipid structures containing 20% GM1, we determined the distribution of GM1 on the membrane and found that GM1 clustering in lipid rafts, thereby supporting the physiological interaction between APOE and GM1. Overall, we find that APOE plays a regulatory role in GM1 transport, and the presence of GM1 on the lipid structures influences this transport process. Our studies introduce a plausible direct link between APOE and AD etiology, wherein APOE regulates GM1, which, in turn, promotes Aβ oligomerization and aggregation.

## INTRODUCTION

AD is a prevalent neurodegenerative disorder that predominantly affects cognitive function, memory, and overall behavior, particularly in the elderly^1–5^. AD is characterized by the presence of Aβ plaques and neurofibrillary tau tangles. Aβ plaques, composed of Aβ proteins, are extracellular deposits that are associated with AD pathology^6–8^. While their precise role in disrupting neuronal function and contributing to synaptic dysfunction and neurodegeneration is still under investigation, they are considered a key pathological feature of AD^7,9^. Intracellular neurofibrillary tangles, primarily composed of hyperphosphorylated tau protein, disrupt the structural integrity of neurons and impair cellular communication^7^. We still do not understand the mechanistic link between the processes leading to Aβ aggregation and the formation of neurofibrillary tau tangles, as well as the role of the protein, APOE, which has been genetically linked to AD. In the brain, APOE is synthesized and secreted by glial cells, particularly astrocytes^10^. Within the human population, three isoforms of APOE exist: APOE4 poses a significant genetic risk factor for late-onset AD, APOE3 represents the most common isoform (77%), and APOE2 serves as a protective isoform in AD (8%)^11,12^. Distinct amino acid sequences characterize these isoforms. APOE2 features cysteine at positions 112 and 158, APOE3 includes a cysteine at 112 and an arginine at 158, while APOE4 carries arginine at both positions 112 and 158^13^.

Despite numerous hypotheses^1,2,14–22^ attempting to unravel the AD molecular etiology, the pathogenesis remains elusive. While many of these hypotheses have focused on the role of Aβ, they often fall short of explaining the risk associated with APOE. The pivotal role of APOE in AD risk underscores the need to explore its potential connection with Aβ pathology. Here, we hypothesize that GM1, a ganglioside abundant in cell membranes, particularly in the nervous system, may serve as the missing link between APOE and Aβ pathology. This hypothesis is driven by two main considerations. First, emerging evidence highlights the significance of membrane composition elements^5,23–25^, such as GM1, in the intricate processes of Aβ aggregation and neuronal protection. GM1, a ganglioside composed of glycosphingolipid and sialic acid, is abundant in cell membranes, particularly in the nervous system^23,26^. It plays diverse roles in cell signaling, recognition, and interaction. GM1 is enriched in lipid rafts and specialized membrane microdomains^27^. It is reported that in the frontal cortex of 10 early-stage AD patients, ganglioside content is comparable to a matched-aged control group, but the percentage of gangliosides in lipid rafts, especially GM1, doubled^26,28^. Alzheimer patients’ platelets exhibited significantly higher GM1 content in lipid rafts than controls, indicating a potential link between GM1 and AD pathology^26^. Other studies^24,37,38,39^, have also underscored the impact of GM1 on Aβ pathologies in AD^37,38,40^. Second, APOE, consisting of 299 amino acids with a molecular mass of approximately 34 kDa, is primarily a critical carrier of cholesterol and other membrane lipids. Following synthesis, APOE undergoes lipidation primarily through adenosine triphosphate-binding cassette transporters A1^29,30^, facilitating the transport of cellular cholesterol and phospholipids to lipid-free APOE within lipoproteins, such as high-density lipoproteins particles^31,32^. Subsequently, APOE transports lipids throughout the brain by interacting with specific receptors on cell surfaces, particularly neurons. These receptors include LDLR, LRPR-related protein 1 (LRP1), VLDL receptor (VLDLR), and APOE receptor 2 (ApoER2)^33^. LDLR, a transmembrane receptor, stands out as a major APOE receptor^34,35^. Given that GM1 is also a lipid, it is plausible that APOE may transport GM1. However, despite prior indications of an age-related rise in GM1 levels in detergent-resistant membranes of APOE4 knock-in mouse brains^3,36^, there is a paucity of research connecting APOE to GM1 in AD pathology. The transportation of GM1 by APOE and whether APOE serves as a lipid carrier for GM1, akin to its role in cholesterol transport, remains an unresolved mystery.

To test our hypothesis, we employ a multifaceted approach, integrating *in vitro*, *in silico*, and cellular studies to meticulously examine the interaction between GM1 and APOE. We assess the binding affinity between APOE and GM1 utilizing MicroScale Thermophoresis (MST) and find that APOE has a higher binding affinity to GM1 than cholesterol, and the binding affinity is affected by the concentration of GM1 on the membrane. We characterize the cellular uptake of GM1 lipid structures under the regulation of APOE and find that APOE facilitates the transport of GM1 to the cell membrane. This transport process is dependent on the cell type, and the expression of APOE receptors in the cell line. In the presence of APOE, GM1 lipid structures compete with cholesterol lipid structures for cellular uptake, exhibiting enhanced cellular uptake compared to cholesterol lipid structures. Using circular dichroism (CD) measurements, we find that GM1 levels on the membrane affect the secondary structure of APOE and increase the binding affinity of APOE-lipoprotein to APOE receptor LDLR. To determine the distribution of GM1 on the membrane, we employ a confocal microscope and discrete molecular dynamics (DMD) simulations to explore the distribution of GM1 on lipid structures and the impact of GM1 levels on the formation of GM1 raft structures. Our findings reveal that a membrane containing a high concentration of GM1 (40%) exhibited instability, whereas the membrane with 20% GM1 maintained its integrity while forming stripe-like clusters on the surface. Our results unveil a potential pathway in which APOE regulates GM1 transport, while simultaneously, GM1 on the membrane influences the transport process. The presence of GM1 on the membrane affects membrane stability and structural integrity, consequently altering the secondary structure of APOE3 and APOE4, and potentially enhancing the binding affinity between APOE and its receptor. The presence of GM1 clusters promotes the cellular uptake of lipid structures under the regulation of APOE3 and APOE4. Through these comprehensive analyses, we establish a definitive link between the distinct modulation of GM1 transport by APOE, one of the mysteries of the of AD pathogenesis.

## RESULTS

### Binding affinity between APOE and lipid structures with different contents

To explore the potential regulatory role of APOE in GM1 transport, we tested the binding affinity between APOE and GM1 and compared the binding affinity of GM1 to cholesterol and other lipids commonly used in membrane lipid structures. Lipid structures were prepared using cholesterol, GM1, sphingomyelin (SM) or L-α-phosphatidylcholine (PC) correspondingly. We find that when sole SM was used, phase separation between SM and the buffer occurred. Consequently, mixed lipid structures of SM and PC were prepared instead of using 100% SM. APOE3 and APOE4 were labeled with RED-NHS. We used MST to determine the binding affinity between various lipid structures and both APOE3 and APOE4. During the formation of lipid structures, lipids can give rise to various configurations, such as lipid bilayers, liposomes, and micelles. Given the presence of internal lipids within the lipid bilayer and the potential encapsulation of some lipids in liposomes, the actual effective concentration of lipids available for binding to APOE was uncertain. Consequently, the Kd here serves as a relative value. Our findings revealed that cholesterol and PC exhibited significantly higher dissociation constant (Kd) values than GM1 and the mixture of SM and PC, for both APOE3 and APOE4. This finding implies that GM1 exhibits superior binding affinity in comparison to cholesterol and PC (Figure 1a, 1b and 1d). Considering APOE’s role in cholesterol transport and GM1’s enhanced affinity to APOE over cholesterol, the results suggest a potential involvement of APOE in GM1 transport. This finding raises the intriguing possibility that APOE regulates the transport of GM1, consequently impacting AD pathology.

**Figure 1.**
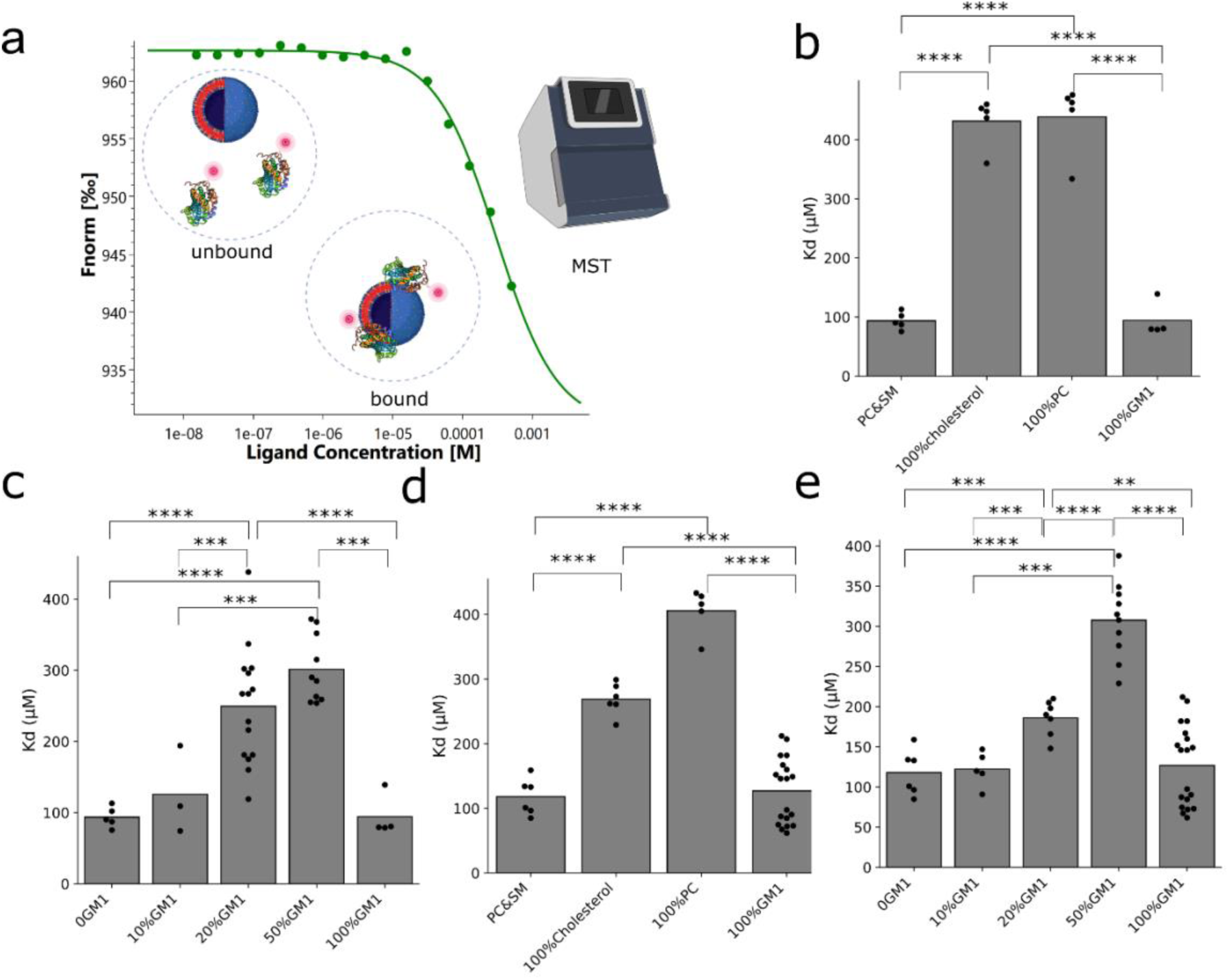
Determination of the interaction affinity between GM1 and APOE. (a) Schematic illustration of determining the binding affinity between lipid structure and APOE using MST. (b) The binding affinity Kd between APOE3 and lipid structures with different compositions. (n>=4) (c) The binding affinity between APOE3 and lipid structures with different GM1 concentrations. (n>=3) (d) The binding affinity between APOE4 and lipid structures with different compositions. (n>=5) (e) The binding affinity between APOE4 and lipid structures with different GM1 concentrations. (n>=5) P-value: ns (0.05 < p <= 1), * (0.01 < p <= 0.05, ** (0.001 < p <= 0.01, *** (0.0001 < p <= 0.001, **** (p <= 0.0001). (a) is created with BioRender.com.

To explore the impact of GM1 levels in membranes on the binding affinity to APOE, we prepared lipid structures with diverse GM1 concentrations (0%, 10%, 20%, 50%, and 100% GM1). We evaluated the binding affinity between lipid structures exhibiting distinct GM1 levels and APOE3 and APOE4, using MST (Figure 1c, 1e). The incorporation of increasing amounts of GM1 in the lipid structures resulted in a diminishing binding affinity. A distinctive pattern emerged when GM1-only lipid structures were used, showing an increasing trend in binding affinity. This observation indicates that the binding affinity between the GM1 membrane and APOE is influenced not only by the concentration of GM1 but also by other factors, such as the size of the lipid structure and membrane deformation resulting from elevated GM1 content. We discuss the effect of the size of lipid structures on membrane deformation by GM1 below.

To explore the impact of lipid structure size on the binding affinity with APOE, we prepared lipid structures with diverse sizes and compositions (Figure S1a-f), evaluating their binding affinity with both APOE3 and APOE4 (Figure S1g, S1h). Our findings reveal that APOE3 has a better binding affinity to small lipid structures formed by GM1 lipids, while for the other two lipid structures, the size of lipid structures does not significantly affect the binding affinity. In contrast, APOE4 demonstrates a heightened binding affinity to larger lipid structures when compared to lipid structures containing cholesterol, PC, and SM. Lipid structures containing GM1 exhibit higher affinity towards both APOE3 and APOE4 compared to those containing cholesterol.

### APOE transports GM1 on cell membranes

To determine whether APOE transports GM1 to the cell membrane, we studied the influence of APOE on the cellular uptake of GM1 lipid structures. Lipid structures containing varying GM1 contents and labeled with DiI were incubated with APOE3 and APOE4 overnight at 37°C, resulting in the APOE-enriched lipoproteins. Subsequently, these lipoproteins were incubated with differentiated PC-12 cells. After removing the excess lipoproteins from the culture medium, we assessed changes in the fluorescence intensity of DiI on the cells. We found a significant increase in cellular uptake when lipid structures contained 20% GM1 in the presence of both APOE3 and APOE4 (Figure S2). To further validate the regulatory effect of APOE on GM1, we determined the cellular uptake of 20% GM1 lipid structures with Human Embryonic Kidney (HEK)-293, U-87 MG, and bEnd.3 cells. U-87 MG and bEnd.3 cell lines are reported to express high levels of APOE receptors^41^. DiI fluorescence intensity analysis demonstrated a significant increase in fluorescence intensity in the presence of APOE3 and APOE4 for U-87 MG and bEnd.3 cells, similar to the observations in differentiated PC-12 cells (Figure 2a, 2b). However, in HEK-293 cells, the fluorescence intensity exhibited no significant change.

**Figure 2.**
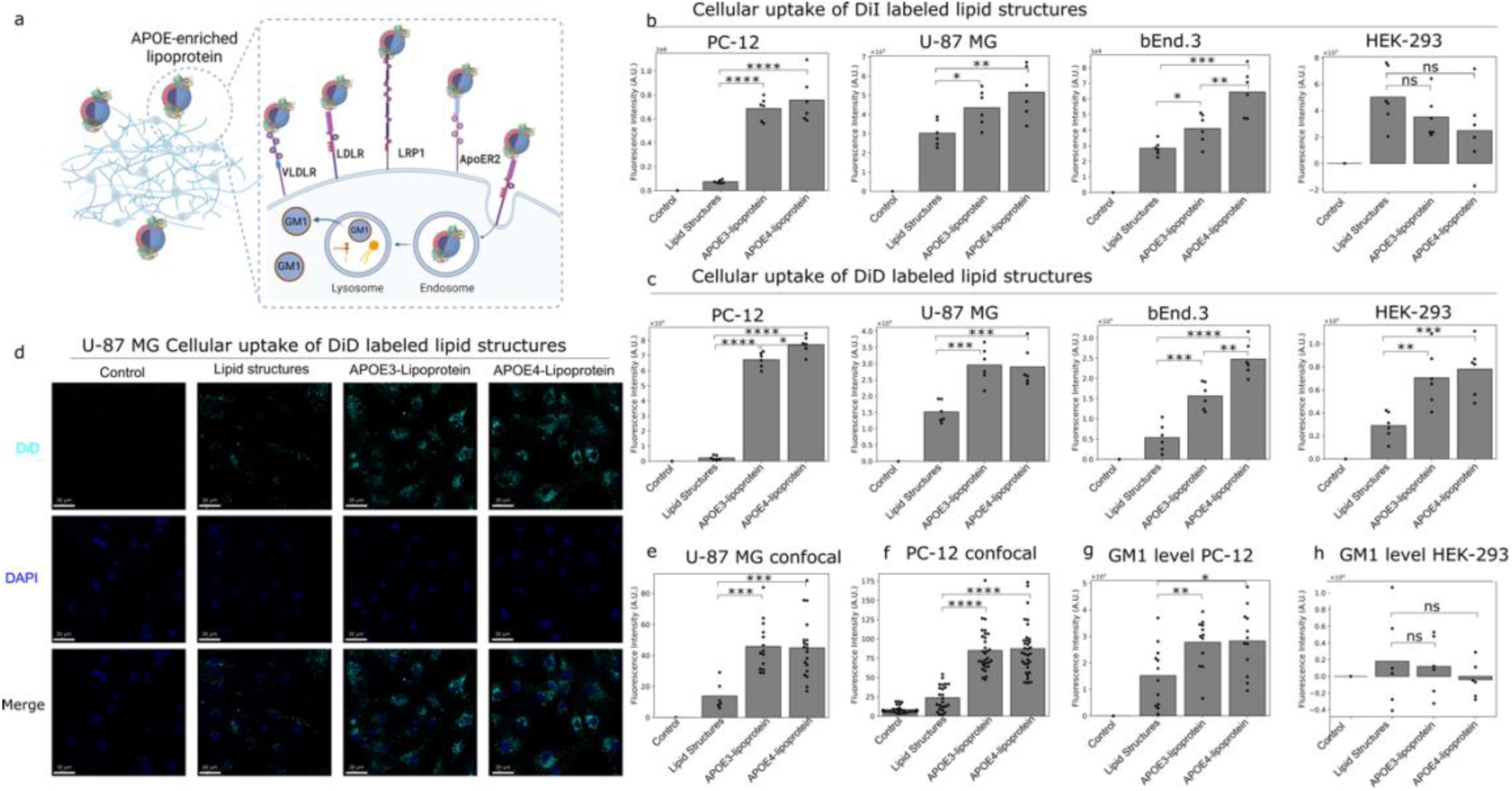
Determination of the cellular uptake of lipid structures. (a) Schematic illustration of cellular uptake of lipid structures. (b) Cellular uptake of DiI labeled lipid structures on differentiated PC-12 cells, U-87 MG, bEnd.3, and HEK-293 cells. (n>=6) (c) Cellular uptake of DiD-labeled lipid structures on differentiated PC-12 cells, U-87 MG, bEnd.3, and HEK-293 cells. (n>=6) (d) Cellular uptake of DiD-labeled lipid structures on U-87 MG determined with confocal microscope. (e) Quantification of the images of U-87 MG cellular uptake of DiD-labeled lipid structures using ImageJ. (f) Quantification of the images of PC-12 cellular uptake of DiD-labeled lipid structures using ImageJ. (g) The changing of GM1 levels on the PC-12 cells after the cellular uptake of GM1 lipid structures. (n>=6). (h) The changing of GM1 levels on the HEK-293 cells after the cellular uptake of GM1 lipid structures. (n>=6). P-value: ns (0.05 < p <= 1), * (0.01 < p <= 0.05, ** (0.001 < p <= 0.01, *** (0.0001 < p <= 0.001, **** (p <= 0.0001). (a) is created with BioRender.com.

We also employed DiD labeling to evaluate the cellular uptake of lipid structures. The results showed that, across all cell types, including differentiated PC-12 cells, U-87 MG, bEnd.3, and HEK-293 cells, the presence of APOE3 and APOE4 enhances the cellular uptake of GM1 lipid structures (Figure 2c). In concert with the DiI-labeled outcomes, differentiated PC-12 cells exhibited the most substantial uptake of lipid structures, and U-87 MG and bEnd.3 cells displayed increased cellular uptake in the presence of APOE3 and APOE4. In contrast to DiI-labeled cellular uptake, HEK-293 cells exhibited significant cellular uptake of DiD-labeled lipid structures in the presence of APOE3 and APOE4. However, when compared to other cell types, HEK-293 cells demonstrated the least amount of GM1 lipid structure uptake. The discrepancy may be attributed to the lower level of cellular uptake in HEK-293 cells, as well as the lower sensitivity of DiI compared to DiD Furthermore, APOE4 exhibited significantly higher cellular uptake than APOE3 in differentiated PC-12 and bEnd.3 cells.

To visually assess the cellular uptake of GM1 lipid structures, we labeled the lipid structures with DiD and utilized confocal microscopy to observe uptake by differentiated PC-12 and U-87 MG cells. Quantification of the images was performed using ImageJ. The results show that, for both differentiated PC-12 and U-87 MG cells, the presence of APOE3 and APOE4 enhances cellular uptake (Figure 2d-2f, Figure S3). Remarkably, APOE4 exhibits significantly higher cellular uptake than APOE3 in differentiated PC-12 cells. These findings are consistent with the previously DiD-labeled fluorescence results. The facilitation of cellular uptake by APOE implies its regulatory role in GM1 transport.

Our findings reveal that the presence of APOE3 and APOE4 regulates the transport of GM1 to the cell membrane. The efficiency of this transport varies depending on the cell type, and the effectiveness of APOE3 and APOE4 differs across different cell lines.

APOE exhibits superior binding affinity to GM1 compared to cholesterol, suggesting that APOE-GM1 lipoprotein would compete more effectively than APOE-cholesterol lipoprotein for cellular uptake. We separately incubated DiD-labeled APOE-cholesterol lipoprotein and DiD-labeled APOE-GM1 lipoprotein with cells and assessed cellular uptake. We observed a higher cellular uptake of DiD-labeled APOE-GM1 lipoprotein (Figure S4a). We then incubated U-87 MG cells with APOE3 and APOE4-enriched cholesterol lipoproteins, as well as a combination of APOE3 and APOE4-enriched cholesterol lipoproteins and APOE3 and APOE4-enriched GM1 lipoproteins and measured the cholesterol levels on the cells. In the presence of APOE-GM1 lipoprotein, the cellular uptake of APOE-cholesterol lipoprotein decreased (Figure S4b). This observation suggests a competitive relationship between APOE-GM1 lipoprotein and APOE-cholesterol lipoprotein during the uptake process, with cells displaying a greater propensity for the uptake of APOE-GM1 lipoproteins, suggesting the broader functions of APOE in lipid metabolism and cellular homeostasis.

To gain a quantitative understanding of the distribution and precise changes in GM1 following the cellular uptake of GM1 lipid structures, we utilize Cholera Toxin Subunit B, Alexa Fluor™ 555 Conjugate (CTSB 555) to label and determine the GM1 content on the cell membrane. This process followed the incubation of APOE-GM1 lipoproteins with differentiated PC-12 cells, coupled with the removal of excess APOE-GM1 lipoproteins from the medium. We find that the presence of both APOE3 and APOE4 results in an increase in GM1 levels on the cell membrane (Figure 2g). This observation implies that both APOE3 and APOE4 actively regulate GM1 transportation by facilitating the cellular uptake of GM1 lipid structures, subsequently influencing the composition of GM1 on the cell membrane. For HEK-293 cells, the presence of APOE3 and APOE4 does not alter the GM1 levels on the cell membrane (Figure 2h). This finding suggests the possibility that APOE plays a role in facilitating the transport and incorporation of GM1 into the cell membrane, and, in turn, may have downstream effects on cellular functions associated with GM1, including intercellular communication, signaling processes, and the promotion of Aβ aggregation. And this process occurs in a cell-type-dependent manner.

### What affects the APOE transportation process?

#### Cell-type dependent expression of APOE receptors

ApoE is predominantly synthesized by glial cells in the brain, while its receptors are abundant in neurons^42^. Increasing evidence suggests that these receptors, widely expressed in most neurons of the central nervous system, play crucial roles in brain development and may significantly impact the pathogenesis of AD^42–44^. As we found, APOE facilitates the transport of GM1 to the cell membrane. Since we found that the cellular uptake of GM1 lipid structures is contingent on the cell type, we analyzed the expression of APOE receptors, including LDLR, LRP1, VLDLR, and ApoER2 on differentiated PC-12 cells, U-87 MG, bEnd.3, and HEK-293 cells. We find that differentiated PC-12 cells express significantly higher levels of LDLR and VLDLR, U-87 MG cells exhibit significantly higher levels of VLDLR and LRP1, and bEnd.3 cells display significantly higher levels of ApoER2 (Figure 3a-d, Figure S5). Differentiated PC-12 cells express the highest overall amount of APOE receptors, whereas HEK-293 cells express the lowest. There is a positive correlation between the expression levels of APOE receptors and the cellular uptake of GM1 lipid structures, with differentiated PC-12 cells showing the highest uptake and HEK-293 cells showing the lowest. This observation implies that the cellular uptake of APOE-delivered GM1 is influenced by the quantity of APOE receptors expressed in the cell.

**Figure 3.**
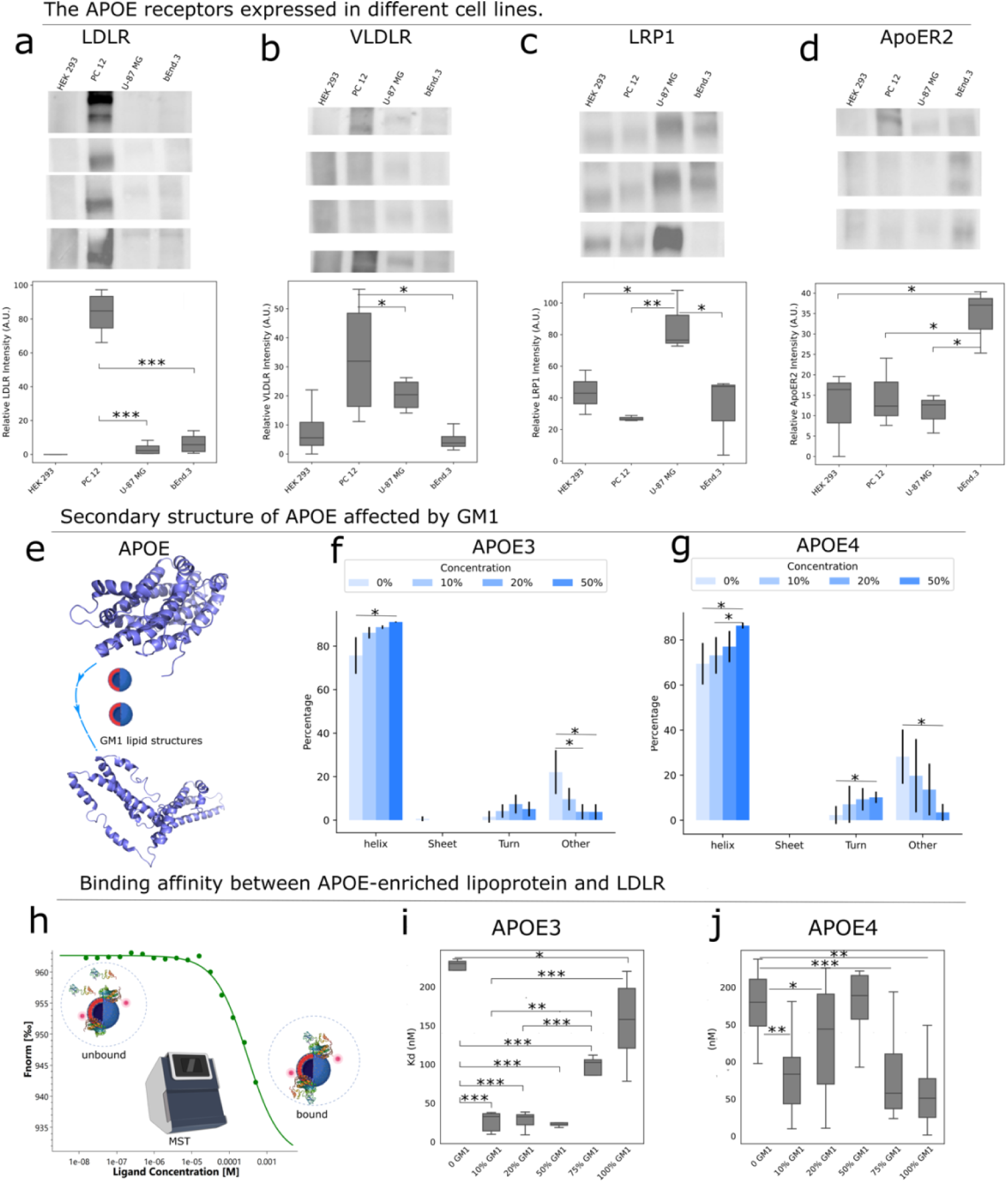
Western blot results of APOE receptors: LDLR, VLDLR, LRP1, ApoER2 on differentiated PC-12 cells, U-87 MG, bEnd.3, and HEK-293 cells. (a) The expression of LDLR on differentiated PC-12 cells, U-87 MG, bEnd.3, and HEK-293 cells (n=4). (b) The expression of VLDLR on differentiated PC-12 cells, U-87 MG, bEnd.3, and HEK-293 cells (n=4). (c) The expression of LRP1 on differentiated PC-12 cells, U-87 MG, bEnd.3, and HEK-293 cells (n=3). (d) The expression of ApoER2 on differentiated PC-12 cells, U-87 MG, bEnd.3, and HEK-293 cells (n=3). (e) Schematic illustration of determining the changes of APOE secondary structures. (f) The CD results of APOE 3 under the effect of different GM1 content on the lipid structures (n>=3). (g) The CD results of APOE 4 under the effect of different GM1 content on the lipid structures (n=3). (h) Schematic illustration of determining the binding affinity of APOE-enriched lipoprotein and APOE receptor LDLR using MST. (i) The binding affinity between APOE3 and lipid structures with different compositions. (n>=4) (j) The binding affinity between APOE4 and lipid structures with different compositions. (n>=3) P-value: ns (0.05 < p <= 1), * (0.01 < p <= 0.05, ** (0.001 < p <= 0.01, *** (0.0001 < p <= 0.001, **** (p <= 0.0001). (e) and (h) are created with BioRender.com.

#### APOE structural changes upon lipid binding

It is well-established that the interaction between APOE and lipids leads to conformational changes in APOE^45,46^. To understand factors influencing the APOE transportation process, we studied whether GM1 binding alters the structural arrangement of APOE, subsequently influencing its recognition by receptors. We characterized the APOE-induced structural changes induced by lipid structures with varying GM1 concentrations by using CD. We incubate GM1 lipid structures with APOE3/APOE4 overnight at 37°C to generate APOE-enriched lipoproteins. We then assess the secondary structure using CD (Figure 3 e-g, Figure S6). We find that, for both APOE3 and APOE4, an increase in GM1 levels within the lipid structures is associated with higher levels of APOE’s helix structure and a reduction in APOE’s other structures. This observation suggests that GM1-induced conformational changes in the secondary structure of APOE.

#### Binding affinity between APOE and LDLR in the presence of GM1

The primary APOE receptors that engage in interactions with APOE, facilitating its role in lipid metabolism and transport, include LRP1, LDLR, APOER2, and VLDLR. LDLR, extensively studied as one of the key APOE receptors, plays a central role in regulating LDL cholesterol levels in the bloodstream^47,48^. APOE’s extensive research history underscores its critical importance in maintaining cholesterol homeostasis^49^. To determine the impact of GM1 content on the binding of APOE to its receptors, we labeled APOE3 and APOE4 with RED-NHS and subsequently incubated them with lipid structures containing varying GM1 levels overnight at 37°C, resulting in the formation of APOE3-lipoprotein and APOE4-lipoprotein. Using MST, we determined the binding affinity between LDLR and APOE3-lipoprotein, as well as APOE4-lipoprotein across different GM1 concentrations. We find that, for both APOE3 and APOE4, in comparison to the control where there are no GM1-containing lipid structures, the presence of GM1 reduces the Kd value (Figure 3h-j), thus suggesting that the presence of GM1 enhances the binding affinity of APOE for LDLR.

In summary, the GM1 levels within the lipid structures induce alterations in the secondary structure of APOE and enhance the binding affinity between APOE and its receptor LDLR. The expression of APOE receptors and the GM1 levels within the lipid structures influence the regulatory transport process mediated by APOE.

### The distribution of GM1 on the membrane

GM1 and other gangliosides, primarily studied for their essential intracellular functions, are most extensively investigated as components of the plasma membrane, particularly in neurons where they serve as the predominant sialoglycans^50^. Within the lipid bilayer, the distinctive structure of GM1 diminishes the fluidity of the plasma membrane and leads to the retention and enrichment of this ganglioside in specific membrane domains, known as lipid rafts^26^. To understand the distribution of GM1 and the influence of elevated GM1 content on membrane morphology and stability, we conducted experiments to evaluate the cellular uptake of lipid structures with varying GM1 concentrations (0%, 10% GM1, 20% GM1, 50% GM1, 100% GM1) in the presence of APOE and employing the PC-12 cell line. A significant improvement in cellular uptake was observed when the GM1 content in the lipid structure was 20%. To understand the reason for this enhanced cellular uptake at 20% GM1, we performed Discrete Molecular Dynamics (DMD)^51–54^ simulations of POPC membrane mixed with 10%, 20%, and 40% GM1 (Figure 4). To reach time scales associated with the lipid/membrane dynamics (the diffusion coefficient of lipids in the membrane, *D* ∼1 μm^2^/s), we adopted a coarse-grained (CG) lipid model with implicit solvent in DMD simulations. GM1 was modeled as a hybrid all-atom head with CG tails because the interactions of the solvent-exposed heads are important in their self-assembly dynamics (Methods). With a constant zero-tension in the x-y plane and a constant temperature of 300 K, the membrane area projected to the plane fluctuated (Figure 4a). For membranes with 10% and 20% GM1, all independent simulations reached their steady states during simulations. With an average area per lipid of ∼0.63 nm^2^, a POPC membrane with 400 lipids per leaflet has an average area of ∼252 nm^2^. The average area of membranes with both 10% and 20% GM1 was larger than that of pure POPC membrane, likely due to the bulky head of GM1 and its net charge of -e, resulting in both steric and electrostatic repulsion between GM1 lipids. Hence, increasing the ratio of GM1 resulted in larger membrane areas. For the membrane with 40% GM1, the initial membrane area in the x-y plane was significantly higher than in the other two cases, but the large membrane area was not sustainable and rapidly decreased after a short period of simulation times in all independent simulations. With a larger surface area but the same amount of lipids, the membrane thickness is expected to be stretched thin and thus the membrane experiences high strain. Examination of snapshots along the simulation trajectories revealed that the membrane underwent rapid deformation and started to crumble (Figure 4b), suggesting that the membrane with such a high amount of GM1 was unstable.

**Figure 4.**
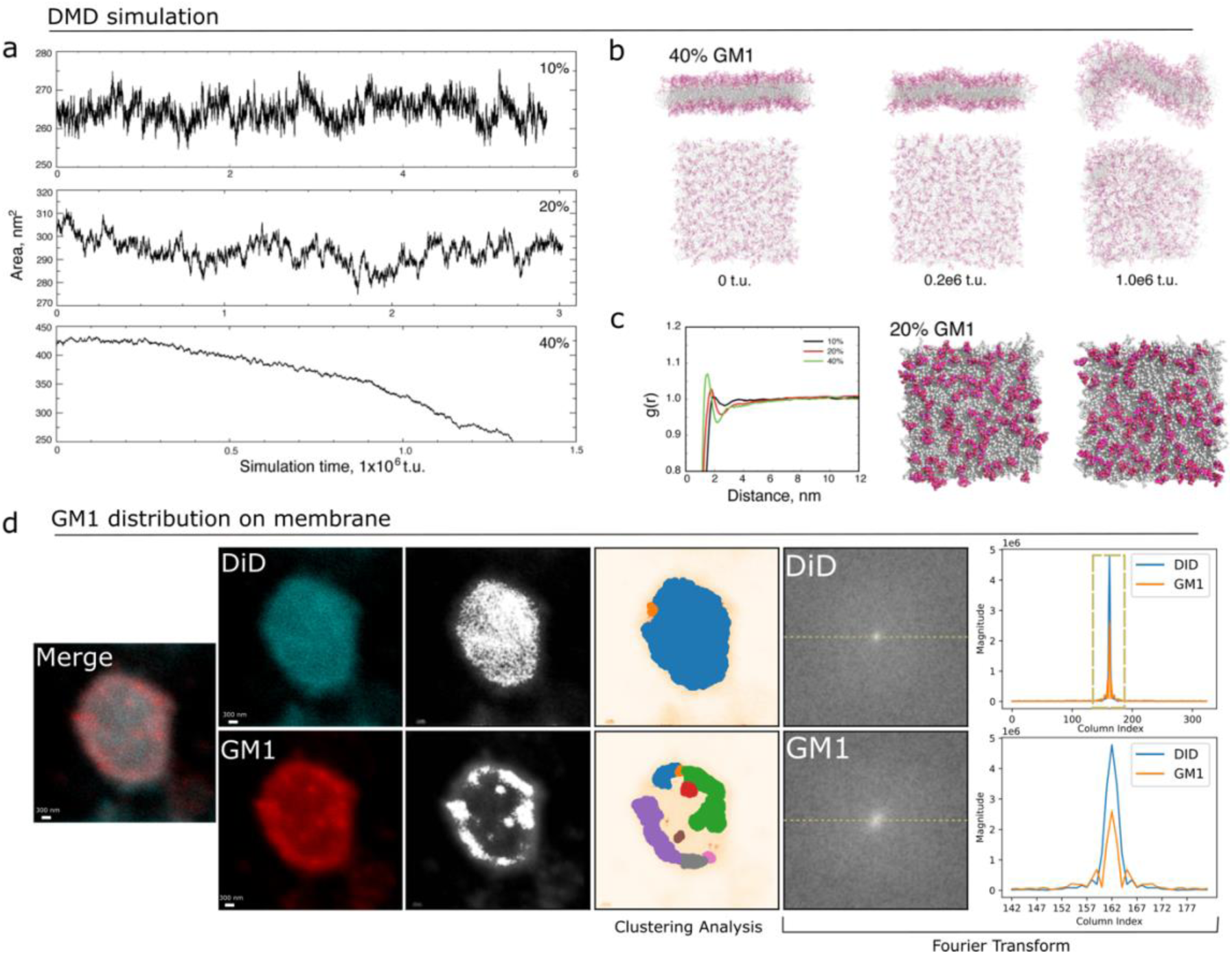
DMD simulations of POPC membranes mixed with different content of GM1: 10%, 20%, and 40%. Ten independent simulations starting with different atomic velocities were performed for each molecular system. (a) Representative trajectories with the time evolution of the membrane area in the x-y plane suggested that membranes with 10% and 20% GM1 were stable and maintained their corresponding structural integrities during the courses of simulations. The DMD time unit (t.u.) corresponds to approximately 50 fs. (b) The membrane with 40% GM1, on the other hand, was unstable and underwent rapid deformation as shown by snapshots at different times along the trajectory. Two views both along the normal of the z-axis or in the x-y plane were shown. Lipid tails were colored gray, and the GM1 heads in red. (c) The pair correlation function, g(r), was obtained for simulations with different ratios. For the case of 40% GM1, the initial metastable phase before the deformation of membranes was used in the calculation. For each GM1 lipid, a central carbon atom in the head was used to represent the lipid. Despite being charged, GM1 had a preference to interact with each other and the g(r) featured a peak at short inter-GM1 distances around 1.6-3 nm. With increasing GM1 content, the pattern became more prominent, suggesting the formation of ordered structures. Two representative snapshots of the membrane with 20% GM1, to the right, illustrated the self-assembly of GM1 lipids into stipe-like clusters. (d) Representative image showcasing the distribution of lipids and GM1 on the lipid structure membrane. All lipids on the lipid structures were labeled with DiD, while GM1 on the lipid structures was labeled with CTSB-555. The merged confocal images of DiD and GM1 (column 1), the individual images of DiD and GM1 (column 2), the grayscale representations (column 3), the pixel clusters (column 4), the 2D Fourier transforms (column 5), and 1D Fourier transforms (column 6) are presented. The 1D Fourier transform represents the central horizontal cross-section in the 2D Fourier transform plot. The lower 1D Fourier transform plot corresponds to the region outlined by the dashed box in the upper 1D Fourier transform plot.

To capture the distribution of GM1 on the membrane, we computed the pair correlation function, g(r), of GM1 lipids (Figure 4c). Here, g(r) denotes the average density of other GM1 lipids within a spherical shell with radii between r and r+dr away from each GM1, normalized by the bulk density of GM1. For the case of 40% GM1, the initial metastable phase before the deformation of membranes was used in the calculation. The pair correlation function featured peaks and valleys at a short distance of around 1.6-3 nm, suggesting the formation of ordered structures. At longer distances, g(r) approached the unity. With increasing GM1 ratios, the peaks and valleys became more distinct with shorter distances. Since the membrane with 40% GM1 was eventually not stable, GM1 with 20% was examined for the structural patterns (Figure 4c). Interestingly, GM1 on the surfaces formed stipe-like clusters. The interaction was likely driven by hydrogen bonds between hydroxyl groups rich in the peripheral of the head group while the single carboxyl group was positioned near the center. Taken together, our DMD simulations suggest that POPC membrane with 20% GM1 features retained membrane integrity while forming stripe-like clusters on the surface, enhancing interactions with APOE.

To elucidate the distribution of GM1 in lipid structures, we implemented a dual-labeling approach, utilizing DiD for all lipids on lipid structures and CTSB-555 for GM1 on lipid structures. Subsequently, we used confocal microscopy to analyze the spatial arrangement of lipids. We converted the confocal images to grayscale, revealing a non-uniform distribution of GM1, in contrast to the more homogeneous distribution observed for other lipids labeled by DiD in the system (Figure 4d). To address potential biases introduced by faint pixel intensities and ensure accurate observations, we applied the DBSCAN algorithm^55^ directly to the pixel clustering in the images. The clustering results indicated that pixels representing DiD were grouped into a single class, while those representing GM1 were distributed across multiple classes. Given that DBSCAN classifies data points based on density, this further suggests that DiD’s distribution is uniform, whereas GM1’s distribution is highly non-uniform. Additionally, we performed a Fourier analysis of the confocal images. The analysis revealed that DiD’s spectrum exhibited a highly intense central peak, typically indicative of a uniform pixel distribution. In contrast, GM1’s spectrum displayed a lower central peak with smaller frequencies around it, indicating a more uneven distribution compared to DiD. Hence, both cluster analysis and Fourier analysis collectively emphasize the highly non-uniform distribution of GM1 on lipid structures, underscoring the importance of accounting for biases associated with faint pixel intensities during visual interpretation. Our experimental observations are in concert with DMD simulations.

In summary, we observed that membranes with an excessive amount of GM1 (40%) were unstable, while those with 20% GM1 maintained integrity and formed stripe-like clusters on the surface. These GM1 clusters may further facilitate the cellular uptake of lipid structures under the regulation of APOE, as discussed above.

## DISCUSSION

The APOE gene, particularly its ε4 allele, has long been recognized as a significant risk factor for late-onset AD^13,43^, yet the precise mechanisms underlying its influence remain elusive despite extensive research. While numerous studies^10,56,57^ have explored APOE’s associations with Aβ clearance^13^, neuroinflammation^6^, and neuronal membrane integrity^58^, conclusive evidence linking APOE4 to AD pathogenesis is still lacking^59^. Our findings contribute to bridging this gap by establishing direct physical and biological connections between APOE and AD pathology. Specifically, APOE’s role in lipid transport, particularly cholesterol^60,61^, is well-documented in human models, with implications for myelin and neuronal membrane repair^62^ and maintenance in both the peripheral and central nervous systems. Moreover, given GM1’s significant impact on AD etiology^23,38^, such as our previous work^39^ and other studies^63–65^ showing that GM1 promotes the aggregation of Aβ, we investigated whether APOE transports GM1 and found that APOE exhibits a higher binding affinity to GM1 than cholesterol, suggesting a competitive relationship between the two lipids for APOE-mediated cellular uptake.

Furthermore, our cellular studies highlight the role of APOE in facilitating GM1 transport on cell membranes, with observations varying across different cell types within the brain. Various cell types within the brain may display diverse levels of APOE-mediated GM1 transport, influenced by the varying expression levels of APOE receptors, which are particularly abundant in neurons^42^. The bEnd.3 cell line, derived from brain tissue and characterized as endothelial cells with an endothelioma phenotype, is commonly employed as a model for studying the blood-brain barrier^41^. Our observations reveal a pronounced cellular uptake of GM1 lipid structures by bEnd.3 cells, suggesting a potential possibility for the transport of GM1 from the brain to the blood and/or vice versa.

Intriguingly, our DMD simulation and cellular studies revealed that membranes containing 20% GM1 maintained integrity and formed stripe-like clusters, promoting the cellular uptake of lipid structures under APOE regulation, although the underlying mechanism remains unclear. Moreover, when examining the effect of GM1 content on APOE and LDLR binding affinity, we observed a larger experimental error in the binding of APOE4 to LDLR compared to APOE3, possibly due to differential GM1 distribution inside and outside of the cluster within lipid structures. This highlights the sensitivity of APOE4 to GM1 distribution and its potential implications for APOE-LDLR binding dynamics.

## METHODS

### Lipid structures and APOE-enriched lipoproteins preparation

We dissolved PC (Sigma-Aldrich), SM (VWR International), cholesterol (Fisher Scientific), and GM1 (Sigma-Aldrich) in a chloroform/methanol (1:1, V:V) mixture at a total concentration of 5 mM. We prepared various lipid structures with different compositions to examine their impact, including 50% PC + 50% SM; 100% PC; 100% cholesterol; 45% PC + 45% SM + 10% GM1; 40% PC + 40% SM + 20% GM1; 25% PC + 25% SM + 50% GM1; 12.5% PC + 12.5% SM + 75% GM1; and 100% GM1. We then dried the mixed solution under vacuum at room temperature and rehydrated the lipid film in PBS buffer (pH 7.4) with a 1 mM lipid concentration at 42°C. After vortexing at least four times for 1 hour, we sonicated the turbid emulsion for 1 hour. Additionally, to assess the effect of lipid structure size, we prepared lipid structures by sonicating the rehydrated suspension for 15 minutes, followed by five cycles of freeze-thawing in liquid nitrogen and a 60°C water bath. Subsequently, we extruded the resulting suspension 16 times using an Avanti extruder with 800 nm pore-size polycarbonate filters for lipid structure filtration. The size distribution of the lipid structures was determined using dynamic light scattering.

We incubated recombinant human APOE3 or APOE4 (Peprotech) with lipid structures (total lipid concentration for the lipid structure: 200 µM) at a concentration of 47.6 µg/mL in a water bath at 37°C overnight to form APOE-enriched lipoproteins.

### Assessment of GM1 lipid structures binding affinity to APOE and APOE-enriched lipid structures interaction with LDLR

We followed the manufacturer’s instructions to label APOE3 and APOE4 using the Monolith protein labeling kit RED-NHS 2nd generation (NanoTemper Technologies). In summary, we mixed 90 µL of 10 µM APOE3 or APOE4 dissolved in PBS with 10 µL of 300 µM RED-NHS 2nd dye by pipetting up and down. Afterward, we incubated the mixture for 30 minutes at room temperature in the dark and removed the excess dye using a column to purify the labeled APOE stock solution. Following the manufacturer’s instructions, we utilized MST to assess the binding affinity of lipid structures with varying GM1 concentrations to APOE3 and APOE4.

We added 10 µL of the diluted APOE solution at a concentration of 20 nM, following the MST manufacturer’s instructions, into each of the tubes numbered 2 to 16 in sequence, excluding tube number 1. We added 20 µL of lipid structures to the first sample tube, then transfered 10 μL to the next tube, mixing it with the APOE solution. We begin with stored lipid structures at a total lipid concentration of 1 mM and repeated this dilution for the remaining tubes. We discarded the 10 μL excess from tube 16. The 16 capillary tubes (NanoTemper Technologies) were inserted into each sample tube to enable sample entry into the capillary, and the capillaries were sequenced and detected using MST. To determine the binding affinity between APOE-lipoprotein and its receptor LDLR (MedChemExpress), we first labeled APOE3 and APOE4 with RED-NHS 2nd generation, as described earlier. Subsequently, we incubated lipid structures with different GM1 concentrations with the labeled APOE at 37°C overnight to produce APOE-enriched lipoprotein. We then used MST to assess the binding affinity between these 20 nM APOE-enriched lipid structures and 250 nM APOE receptor, LDLR. We repeated each test at least three times.

### Uptake of GM1 lipid structures by cells

We obtained PC-12 cells from the American Type Culture Collection (ATCC) and culture and differentiated them according to previously published methods^39^. We obtained HEK-293 cells from Jong Yun Lab (Penn State College of Medicine) and cultured them in DMEM (VWR International) supplemented with 10% fetal bovine serum (Fisher Scientific), 1% Non-Essential Amino Acids (Fisher Scientific), and 1% Penicillin/Streptomycin (Fisher Scientific) at 37°C with 5% CO_2_. Then, we seeded HEK-293 cells in 96-well plates at 5000 cells/well. Similarly, we obtained bEnd.3 cells from the ATCC and cultured in DMEM supplemented with 10% fetal bovine serum and 1% Penicillin/Streptomycin at 37°C with 5% CO_2_. Next, we seeded bEnd.3 cells in 96-well plates at 5000 cells/well. We obtained U-87 MG cells from Jeffrey Neighbors lab (Penn State College of Medicine) and cultured them in Eagle’s Minimum Essential Medium (ATCC) supplemented with 10% fetal bovine serum and 1% Penicillin/Streptomycin at 37°C with 5% CO_2_. Subsequently, we seeded U-87 MG cells in 96-well plates at 5000 cells/well.

We labeled 500 µL of 1 mM lipid structures with different contents by adding 10 µL of 1 mM DiI Stain (1,1’-dioctadecyl-3,3,3’,3’ tetramethylindocarbocyanine perchlorate) (Fisher Scientific) or 10 µL of 1 mg/mL Invitrogen™ DiD oil (DiD, Fisher Scientific) to the lipid solution during preparation. We incubated DiI or DiD-labeled 0.5 mM lipid structures with 47.6 µg/mL APOE3 or APOE4 at 37°C overnight to form APOE-enriched lipoproteins. To investigate the impact of GM1 content on cellular uptake, we incubated differentiated PC-12 cells with various 15 µM lipid structures (50% PC + 50% SM; 45% PC + 45% SM + 10% GM1; 40% PC + 40% SM + 20% GM1; 25% PC + 25% SM + 50% GM1; 12.5% PC + 12.5% SM + 75% GM1; and 100% GM1), and the correspondingly APOE3 and APOE4 (3.66 µg/mL)-enriched lipoproteins for 4 hours, using seeded cells at a density of 2000 cells per well. To characterize the cellular uptake of lipid structures in different cell lines, we added APOE3 or APOE4-enriched lipoproteins (40% PC + 40% SM + 20% GM1) to differentiate PC-12, bEnd.3, U-87 MG, and HEK-293 cells and incubated them for 4 hours, using seeded cells at a density of 2000 cells per well. We added equivalent quantities of lipid structures or PBS to the cells as controls for comparison with APOE-enriched lipoproteins. We removed excess DiI and DiD-labeled APOE-enriched lipoproteins from the medium by thoroughly washing with PBS three times. We measured the fluorescence intensity of DiI and DiD using SpectraMax i3 to elucidate the uptake of APOE-enriched lipoproteins. For DiI, we used the excitation wavelength of 522 nm, and the emission wavelength of 560 nm. For DiD, we used the excitation wavelength of 644 nm, and the emission wavelength of 670 nm. Each sample consisted of 6 wells.

To assess the cellular uptake of cholesterol, we incubated U-87 MG cells with DiD-labeled APOE-enriched cholesterol lipoproteins (40% PC + 40% SM + 20% cholesterol) and DiD-labeled APOE-enriched GM1 lipoproteins (40% PC + 40% SM + 20% GM1) overnight and remove the excess lipoproteins to determine the fluorescence intensity. To investigate the potential competition between GM1 and cholesterol lipid structures in cellular uptake, we incubated U-87 MG cells with APOE-enriched cholesterol lipoproteins (40% PC + 40% SM + 20% cholesterol), as well as a combination of APOE3 and APOE4 (3.66 µg/mL)-enriched cholesterol lipoproteins (40% PC + 40% SM + 20% cholesterol) and APOE3 and APOE4 (3.66 µg/mL)-enriched GM1 lipoproteins (40% PC + 40% SM + 20% GM1) overnight. We determined the changing of cholesterol level on the cell membrane using Amplex™ Red Cholesterol Assay Kit (Thermo Scientific) according to the manufacturer’s instructions.

To characterize the changes in GM1 levels on the cell membrane, we incubated differentiated PC-12 cells in a 96-well plate with APOE (3.66 µg/mL)-enriched GM1 lipoproteins. After washing to eliminate surplus APOE-enriched GM1 lipoproteins, we added 100 µL of 1 µg/mL CTSB to each well and incubated with cells for 30 minutes at room temperature to label GM1 on the cell membrane. Then, we remove the excess CTSB by washing it with PBS three times. We measured the fluorescence intensity of CTSB on the cell membrane to characterize the changes in GM1 levels using SpectraMax i3 with an excitation wavelength of 555 nm and an emission wavelength of 575 nm.

We coated the cell culture chamber slide with poly-L-Lysine (Bio-Techne) at 37°C for 30 min and seeded cells at 20,000 cells/well. After incubating cells with DiI and DiD labeled APOE (3.66 µg/mL)-enriched lipoproteins overnight, we removed the excess APOE-enriched lipoproteins in the medium by washing with PBS three times. We fixed the cells with 600 µL of 4% paraformaldehyde solution (VWR International) in PBS for 15 min and washed them with PBS three times. We labeled GM1 on the cell membrane with 300 µL of 1 µg/mL CTSB for 30 min at room temperature, followed by washing with PBS three times. We performed nuclear labeling using 600 µL of 2 µg/mL DAPI (Sigma-Aldrich) for 10 min, and we washed the cells with PBS three times. We used confocal microscopy (LEICA SP8 STED 3X) to determine the cellular uptake of the lipid structures and analyzed fluorescence intensity using ImageJ.

### Cell Lysis and APOE receptor protein quantification

After culturing the differentiated PC-12, bEnd.3, U87-MG, and HEK-293 cells, we added cold RIPA buffer with Pierce™ Protease Inhibitor (1 tablet/10 mL, Thermo Scientific) to the cells and incubated them at 4°C for 15 minutes. Subsequently, we scraped and transferred the cell lysate to 1.5 mL tubes and used sonication to break up any remaining cells with one pulse at 45% power on ice. After centrifuging the lysates at 12,000 rpm for 15 minutes at 4°C, we discarded the pellet. The concentration of total protein was quantified using a BCA kit. To do this, we diluted the obtained total protein from cells 10 times with 10 µL in duplicate wells, added 200 µL of BCA reagent (mix 50:1 A:B), and incubated it for 30 minutes at 37°C. The absorption was then determined at 562 nm using SpectraMax i3.

We employed Western Blot to quantify the levels of APOE receptors, including LRP1, LDLR, APOER2, and VLDLR in different cell lines. The loaded samples were normalized according to BCA quantification results, and the samples were heated at 95°C for 10 min. A 8 % polyacrylamide gel was used for the Western Blot. We used primary antibodies included recombinant anti-LDL receptor antibody (diluted 1:500, Abcam), anti-LRP1 antibody produced in rabbit (diluted 1:1000, Sigma-Aldrich), anti-ApoER2 antibody (diluted 1:1000, Abcam), anti-VLDLR antibody produced in rabbit (diluted 1:1000, Sigma-Aldrich), and control GAPDH antibody (diluted 1:1000, Cell Signaling Technology). We used anti-rabbit HRP (diluted 1:1000, VWR International) as the secondary antibody. We quantified bands using ImageJ.

### Secondary structure

We incubated APOE3 or APOE4 (47.6 µg/mL) in the presence of 1 mM lipid structures with different GM1 contents (50% PC + 50% SM; 45% PC + 45% SM + 10% GM1; 40% PC + 40% SM + 20% GM1; 25% PC + 25% SM + 50% GM1) overnight in a water bath at 37°C. We determined the secondary structure characteristics of APOE3 and APOE4 using a Jasco J-1500 circular dichroism spectrophotometer, using the relative lipid structures solution as the control. We placed samples in 0.1-mm path-length (200 μL) cuvettes, and we recorded the spectra in the range of 185-240 nm with a 0.5 nm data pitch and scanning speed of 50 nm/min.

### The distribution of GM1 on the membrane

We labeled all lipids on the lipid structures membrane using DiD, as mentioned earlier. Subsequently, we labeled GM1 on the 10% GM1 lipid structures by incubating 50 µL of 100 µg/mL CTSB with 100 µL of lipid structures for 30 minutes in the dark and removed the excess CTSB by washing with PBS using a 300 kDa MWCO centrifugal filter tube. We coated the microscope slides with poly-L-Lysine, and we determined the distribution of GM1 on the lipid structures membrane using a confocal microscope (LEICA SP8 STED 3X).

### Discrete molecular dynamics (DMD) simulation

DMD is a rapid and versatile molecular dynamics algorithm. In this study, we modeled the membrane by coarse-grained (CG) lipids with implicit solvent. We maintained a constant pressure with zero-tension in the CG membrane simulations using a Berendson barostat. Briefly, each POPC (1-palmitoyl-2-oleoyl-sn-glycero-3-phosphocholine) lipid was represented by 15 CG beads including one choline, one phosphate, and one glycerol bead in the head, as well as two ester beads connecting ten hydrophobic beads forming two linear tails. The intra- and inter-CG lipid interactions were self-consistently derived using an iterative Boltzmann inversion method to reproduce all the pair correlation functions observed in equilibrium all-atom molecular dynamics simulations with explicit solvent at room temperature and ambient pressure. GM1, on the other hand, had a hybrid representation in our simulation. The head of GM1 lipid had all-atom representation, while two tails of GM1 were coarse-grained the same as the CG POPC lipids. The interactions of the all-atom GM1 heads were modeled by the MedusaScore force field, an extension of the Medusa force field for small-molecule ligands that were able to accurately capture protein-ligand interactions.

Initial structures of three membrane systems with 10%, 20%, and 40% GM1 were generated by CHARMM-GUI. Each of the membranes was composed of 800 lipids with 400 in each leaflet. For each molecular system, 10 independent simulations with different initial atomic velocities were performed. With more hybrid GM1 lipids, more calculations were required to reach the same simulations. With 10% GM1, each independent simulation reached ∼5.5×10^6^ time units (t.u.), corresponding to ∼270 ns, and the accumulative time was about ∼2.7 us. The time unit is equivalent to 50 fs. With 20% GM1, each independent simulation lasted ∼3×10^6^ t.u., or ∼150 ns, and the accumulative time was ∼1.5 us. For the system with the most GM1, each independent simulation lasted ∼1.4×10^6^ t.u., or ∼70 ns, and the accumulative time was ∼0.7 μs. The equilibration of the membrane system in the constant tension DMD simulations could be evaluated from the time evolution of the membrane area in the x-y plane (e.g., Fig. 4).

## ACKNOWLEDGEMENTS

We acknowledge support from the National Institutes for Health 1R35 GM134864, the Huck Institutes of the Life Sciences, and the Passan Foundation. The content is solely the responsibility of the authors and does not necessarily represent the official views of the NIH. We acknowledge Jeffrey Neighbors and Jong Yun for providing us with the cells. We also acknowledge the help from Brianna L. Hnath, and Martin V. Dokholyan with the experiments.

