## Supplemental Figure for "APOE regulates the transport of GM1"

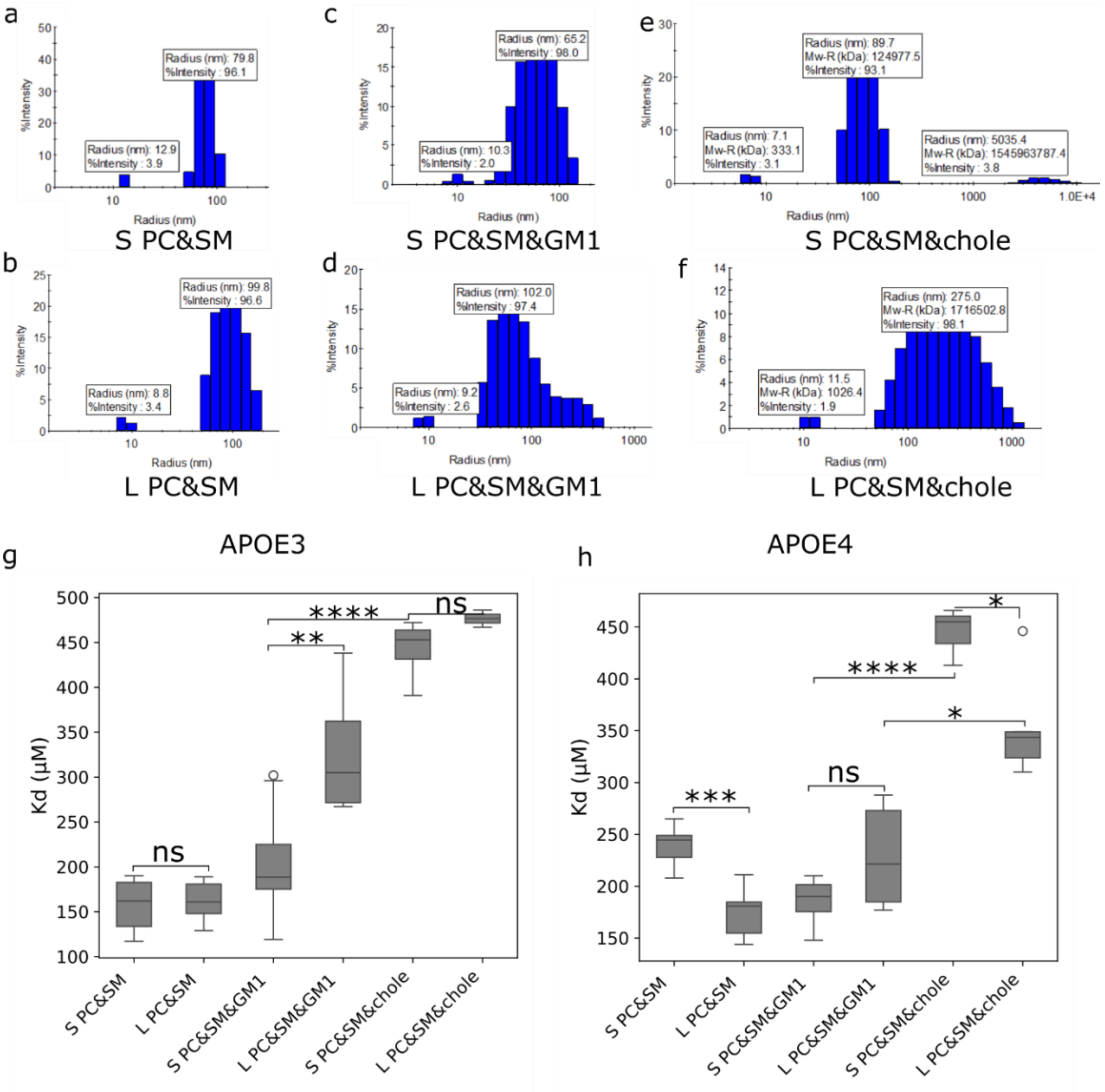


**Figure S1. Determination of the effect of lipid structure sizes and lipid species' composition on the binding affinity Kd between lipid structures and APOE.** (a) Size distribution of the small SM and PC lipid structures (S PC&SM, 50% SM + 50% PC) determined with dynamic light scattering. (b) Size distribution of the large SM and PC lipid structures (L PC&SM,50% SM+50% PC) determined with dynamic light scattering. The lipid structures are prepared using 800 nm pore size polycarbonate filters. (c) Size distribution of the small SM, PC and GM1 lipid structures (S PC&SM&GM1, 40% SM+40% PC+20% GM1) determined with dynamic light scattering. (d) Size distribution of the large SM, PC and GM1 lipid structures (L PC&SM&GM1, 40% SM+40% PC+20% GM1) determined with dynamic light scattering. The lipid structures are prepared using 800 nm pore size polycarbonate filters. (e) Size distribution of the small SM, PC and cholesterol lipid structures (S PC&SM&chol, 40% SM+40% PC+20% cholesterol) determined with dynamic light scattering. (f) Size distribution of the large SM, PC and cholesterol lipid structures (L PC&SM&chol, 40% SM+40% PC+20% cholesterol) determined with dynamic light scattering. The lipid structures are prepared using 800 nm pore size polycarbonate filters. (g) The binding affinity between lipid structures with different compositions and APOE3. (n>=3) (h) The binding affinity between lipid structures with different GM1 concentration and APOE4. (n>=3) P-value: ns (0.05 < p <= 1), * (0.01 < p <= 0.05, ** (0.001 < p <= 0.01, *** (0.0001 < p <= 0.001, **** (p <= 0.0001).

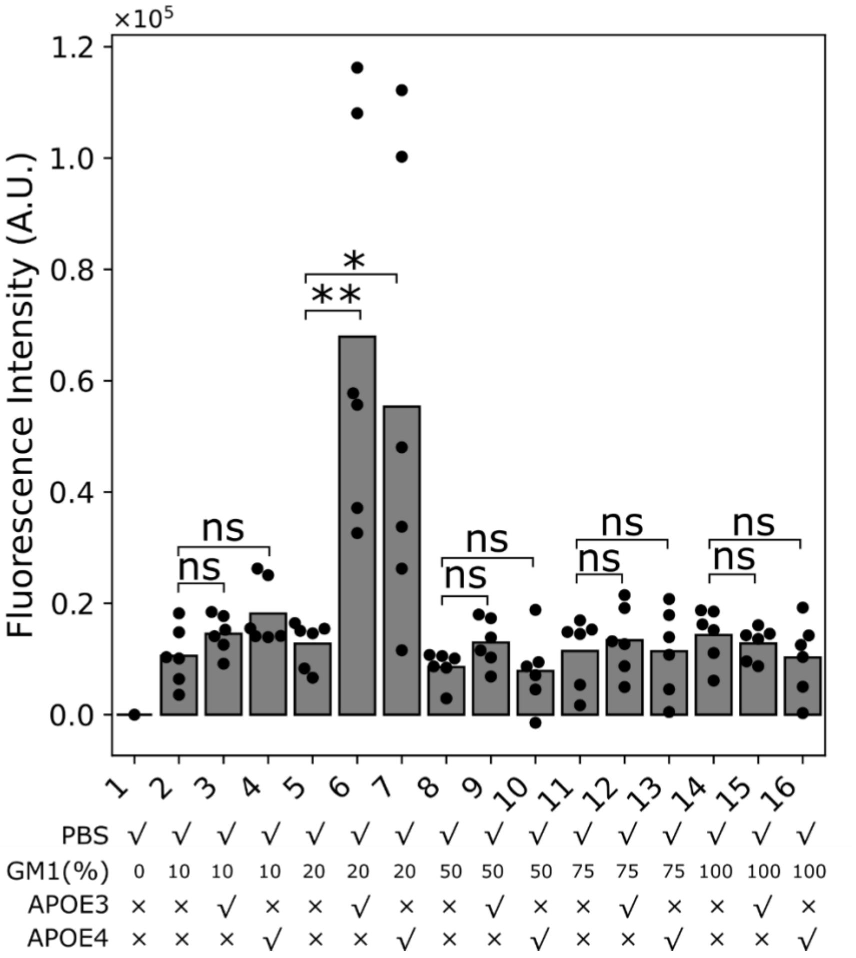


**Figure S2. Determination of PC-12 cellular uptake of lipid structures with varying GM1 concentrations.** We seeded PC-12 cells in 96-well plates at a density of 2000 cells/well and differentiated PC-12 for 1 week. Subsequently, we incubated DiI-labeled lipid structures with varying GM1 concentrations, along with corresponding APOE3 and APOE4-enriched lipoproteins with cells for 4 hours, and then we determined the cellular uptake. (n>=6) P-value: ns (0.05 < p <= 1), * (0.01 < p <= 0.05, ** (0.001 < p <= 0.01, *** (0.0001 < p <= 0.001, **** (p <= 0.0001).

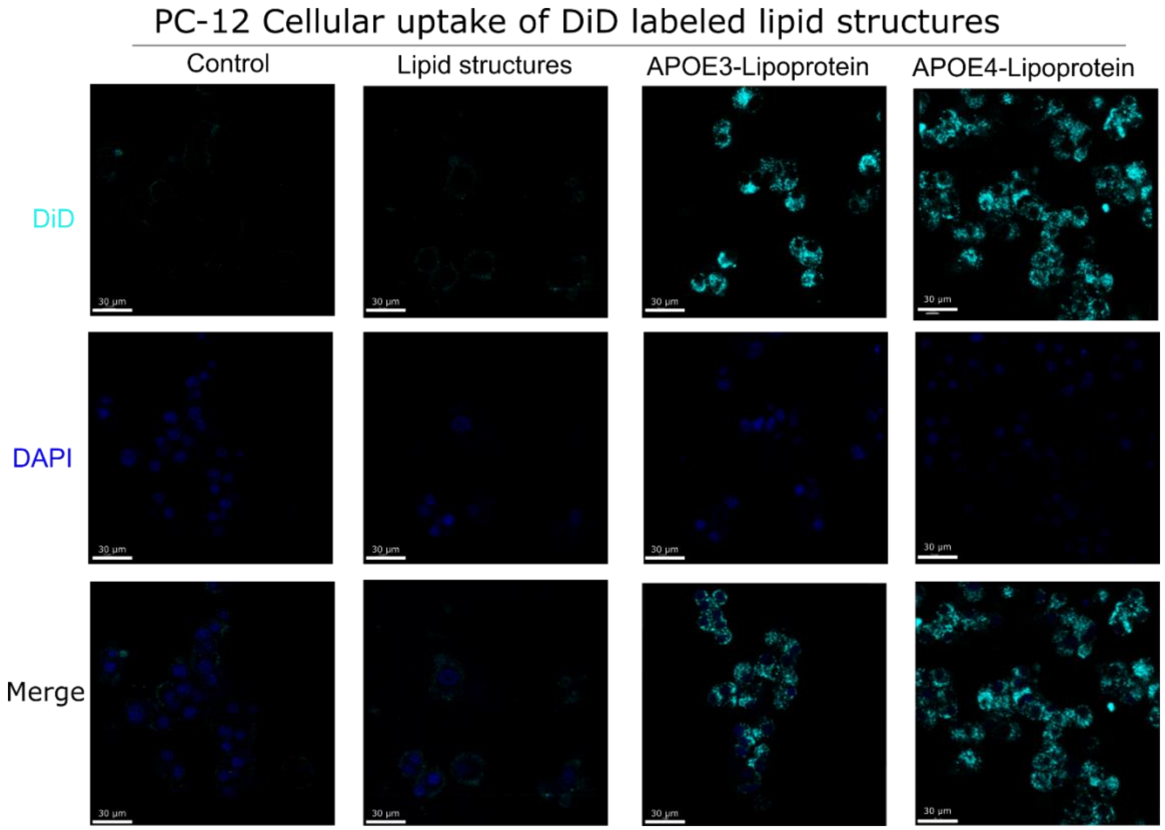


**Figure S3.** Cellular uptake of DiD-labeled lipid structures on PC-12 cells determined with confocal microscope.

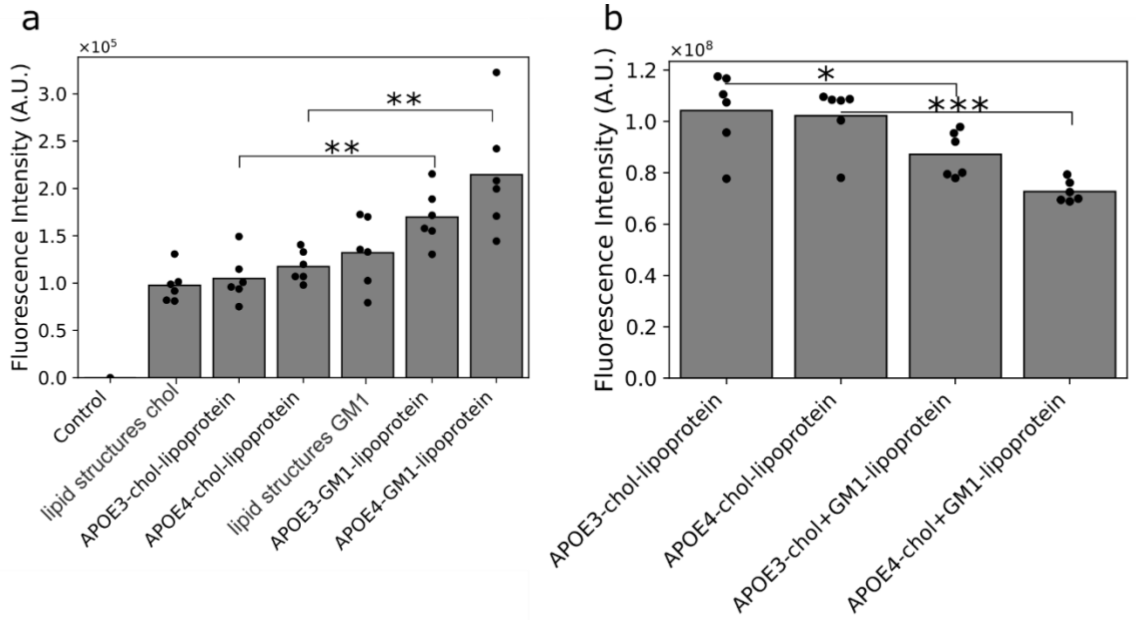


**Figure S4. Determination of cellular uptake of cholesterol and GM1 lipid structures.** (a) Cellular uptake of DiD-labeled cholesterol and GM1 lipid structures on U-87 MG cells. We incubated APOE3 and APOE4 enriched DiD-labeled cholesterol lipoproteins and DiD-labeled

GM1 lipoproteins with the U-87 MG cells overnight. Then, we determined the cellular uptake. (n>=6) (b) Changes in cholesterol levels in U-87 MG cells after cellular uptake of APOE3 and APOE4-enriched cholesterol lipoproteins with U-87 MG cells and cellular uptake of a mixture of APOE3 and APOE4 enriched cholesterol lipoproteins and APOE3 and APOE4-enriched GM1 lipoproteins with U-87 MG cells. We incubated U-87 MG cells with APOE3 and APOE4-enriched cholesterol lipoproteins, as well as a combination of APOE3 and APOE4-enriched cholesterol lipoproteins and APOE3 and APOE4-enriched GM1 lipoproteins overnight and then measured the cholesterol levels on U-87 MG cells. (n>=6). P-value: ns (0.05 < p <= 1), * (0.01 < p <= 0.05, ** (0.001 < p <= 0.01, *** (0.0001 < p <= 0.001, **** (p <= 0.0001).

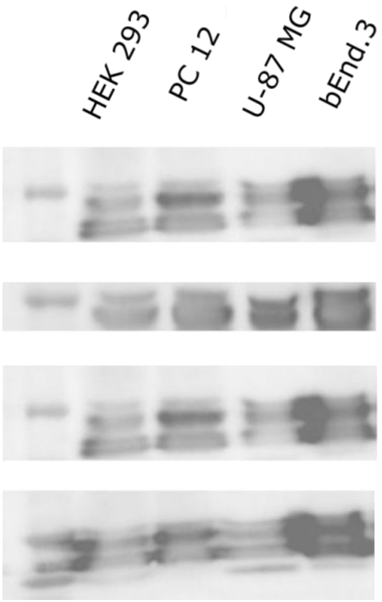


**Figure S5.** Western blot analysis of GAPDH control in various cell lines.

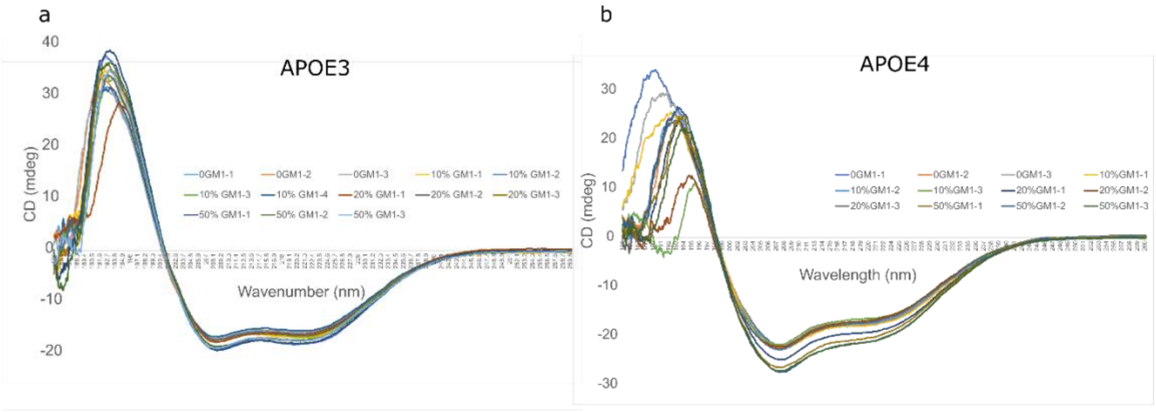


**Figure S6.** Secondary structures of APOE3 and APOE4 under the effect of varying GM1 concentrations within the lipid structures. (a) CD results of APOE 3 with different GM1 content in the lipid structures. (b) CD results of APOE 4 with different GM1 content in the lipid structures.
